# DHFR metabolic activity controls neurogenic transitions in the developing Human and mouse neocortex

**DOI:** 10.1101/2022.06.22.497156

**Authors:** Sulov Saha, Thomas Jungas, David Ohayon, Christophe Audouard, Tao Ye, Mohamad-Ali Fawal, Alice Davy

## Abstract

One-carbon/folate (1C) metabolism supplies methyl groups required for DNA and histone methylation, and is involved in the maintenance of self-renewal in stem cells. Dihydrofolate reductase (DHFR), a key enzyme in 1C metabolism, is highly expressed in Human and mouse neural progenitors at the early stages of neocortical development. Here, we investigated the role of DHFR in the developing neocortex and report that reducing its activity in Human cerebral organoids and mouse embryonic neocortex accelerates indirect neurogenesis, a hallmark of mammalian brain evolution, thereby affecting neuronal composition of the neocortex. Further, we show that decreasing DHFR activity in neural progenitors leads to a reduction in One-carbon/folate metabolites and correlates with modifications of H3K4me3 methylation. Our findings reveal an unanticipated role for DHFR in controlling specific steps of neocortex development and indicate that variations in 1C metabolic cues impact cell fate transitions.

## INTRODUCTION

One-carbon metabolism is composed of two intertwined cycles, the folate cycle whose main output is purine synthesis which is necessary for DNA replication and cell proliferation, and the methionine cycle whose main output is methyl group synthesis which are used for methylation reactions and deposition of epigenetic marks (Clare et al., 2018; Ducker and Rabinowitz, 2017). While the folate cycle has been extensively studied for its role in cell proliferation, recent studies revealed a role for the methionine cycle in self-renewal of stem cells through the regulation of histone methylation (Fawal et al., 2018, 2021; Shiraki et al., 2014; Shyh-Chang et al., 2013). Specifically, it has been shown that perturbation of the methionine cycle leads to a decrease in H3K4 trimethylation, which in turn impacts gene expression (Fawal et al., 2018; Mentch et al., 2015; Shiraki et al., 2014; Shyh-Chang et al., 2013). Dihydrofolate reductase (DHFR), an enzyme positioned upstream of the two cycles, is critical for converting dietary folate into Tetrahydrofolate (THF), and its downstream folate species are required for both DNA synthesis and methylation reactions. DHFR is classified as a housekeeping gene expressed in all cycling cells whose expression varies proportionally to cell growth (Feder et al., 1989). Yet, Human genetic studies showed that DHFR deficiency leads to hematological and neurological phenotypes in Humans (Banka et al., 2011; Cario et al., 2011) indicating that DHFR has tissue-specific functions. Furthermore, a recent single-cell gene expression analysis revealed that *DHFR* mRNA is dynamically expressed in Human and mouse neural progenitors over the course of neocortex development (Klingler et al., 2021) suggesting that it could play specific roles in this developmental process.

In Humans and other mammals, neocortex development begins with neuroepithelial cells as the founder population during embryonic growth. Neuroepithelial cells expand their population before transitioning into apical progenitors residing at the ventricular zone (Noctor et al., 2001). These progenitors express the transcription factor PAX6 (Götz et al., 1998) and give rise to glutamatergic projection neurons populating the six layers of the neocortex, numbered I to VI, in a sequential ‘inside-out’ manner except for early-born uppermost layer I neurons. In the mouse neocortex, at the onset of neurogenesis, a fraction of PAX6^+^ apical progenitors generate early-born deep layer (V-VI) neurons by direct neurogenesis (Cárdenas et al., 2018). During the mid-late stages of neurogenesis, PAX6^+^ apical progenitors generate neurons mostly by indirect neurogenesis which involves the production of intermediate progenitors expressing the transcription factor TBR2 (Haubensak et al., 2004; Miyata et al., 2004; Noctor et al., 2004; Sessa et al., 2008). While TBR2^+^ basal progenitors in the subventricular zone (SVZ) symmetrically divide to generate neurons of all layers, at later stages they mainly give rise to late-born upper layer (II-IV) neurons. In Humans, the number of TBR2^+^ basal progenitors and other types of intermediate progenitors is massively expanded (Hansen et al., 2010) and neurogenesis is mostly indirect although evidence of direct neurogenesis at early developmental stages has been reported for the production of deep layer neurons (Eze et al., 2021). Newborn neurons subsequently migrate radially towards the cortical plate (CP) and express specific factors which control their final positioning and axonal targeting to establish circuit connections, such as TBR1 for layer VI neurons; CTIP2 for layer V neurons, or SATB2 for layer II-IV neurons (Molyneaux et al., 2007).

Here, to address whether DHFR plays a role in specific steps of neocortex development, we reduced DHFR activity both in Human and mouse models, with respectively *in vitro* and *in vivo* approaches. First, we report that DHFR pharmacological inhibition at the initial stage of Human cerebral organoid (HCO) development leads to a depletion in PAX6^+^ apical progenitors and overproduction of TBR2^+^ basal progenitors resulting in accelerated generation of CTIP2^+^ early born neurons and SATB2^+^ late born neurons. Second, we generated haploinsufficient *Dhfr^+/Δ^* mice and observed a loss of PAX6^+^ apical progenitors and CTIP2^+^ early born neurons associated to an increase in TBR2^+^ basal progenitors and SATB2^+^ late born neurons. Both findings suggest that DHFR deficiency prematurely initiates indirect neurogenesis and has a functional impact on the generation of neuron subtypes. Mechanistically, we find that these changes in neuronal subtype generation correlate with decreased steady-state levels of THF and S-Adenosyl methionine (SAM) metabolites and with a global decrease in H3K4 trimethylation (H3K4me3) marks. Genome-wide analyses revealed changes in this histone methylation mark at genes specific for neuronal subtypes.

## RESULTS

### DHFR activity delays neuronal production in Human cerebral organoids

To assess the role of DHFR activity in Human developing brain, we generated HCO with the Lancaster protocol (Lancaster and Knoblich, 2014) and exposed HCO to methotrexate (MTX), a powerful DHFR inhibitor. *DHFR* mRNA expression is highest at early stages of HCO development and is subsequently down-regulated (Figure S1A), we thus treated 7 days *in vitro* (div7) neural aggregates with MTX for a duration of 3 days, with subsequent steps of HCO maturation done in absence of MTX (Figure 1A). Measurement of DHFR activity in control HCO showed a developmental reduction in DHFR activity (Figure 1B), in line with the single-cell mRNA expression data (Figure S1A). Acute 3-day MTX treatment transiently inhibited DHFR since DHFR activity was undetectable in MTX-treated div10 HCO but was recovered at div20 (Figure 1B). Macroscopic observation of HCO over a 60-day time period revealed that MTX treatment stunted HCO overall growth (Figure S1B), yet, in these smaller HCO, the neocortical thickness was increased at div40 and div60 (Figure S1C). In contrast, the thickness of the VZ, which reflects the size of the progenitor pool, remained unchanged at div40 and div60 (Figure S1D), suggesting that the increase in total thickness at these stages might be due solely to the enlargement of the CP in MTX-treated HCO. To further characterize the consequence of transient DHFR inhibition on Human neocortex development, we examined specific progenitor and neuronal populations at the different stages of HCO development and maturation. At div20, MTX treatment led to a significant increase in the fraction of PAX6^+^ apical progenitors and a decrease in the fraction of immature TBR1^+^ neurons (Figures 1C, E). In addition, the fraction of Phospho-histone H3 (pH3)-positive cells was increased in MTX-treated HCO (Figures 1D, E). Altogether these results indicate that inhibition of DHFR leads to an initial reduction and/or a delay in neuronal production at early stages of HCO development. In contrast, at div40, we detected a reduction in the fraction of PAX6^+^ apical progenitors associated with an overproduction of TBR2^+^ basal progenitors and early-born CTIP2^+^ deep layer neurons (Figures 1F-H) and this altered ratio was also present at div60 as we detected a persistent deficit in PAX6^+^ apical progenitors associated with an increase in young TBR1^+^ neurons and early-born CTIP2^+^ neurons (Figures 1I, K). Furthermore, at div60, late-born SATB2^+^ neurons could be observed in MTX-treated HCO while still rare in control samples (Figures 1J, K), suggesting that DHFR inhibition leads to an accelerated neurogenesis. Altogether, these findings indicate that DHFR activity is required in apical progenitors to delay neurogenesis and suggest that transient inhibition of DHFR in Human neural progenitors has long-term impact on the neurogenic pace.

**Figure 1.**
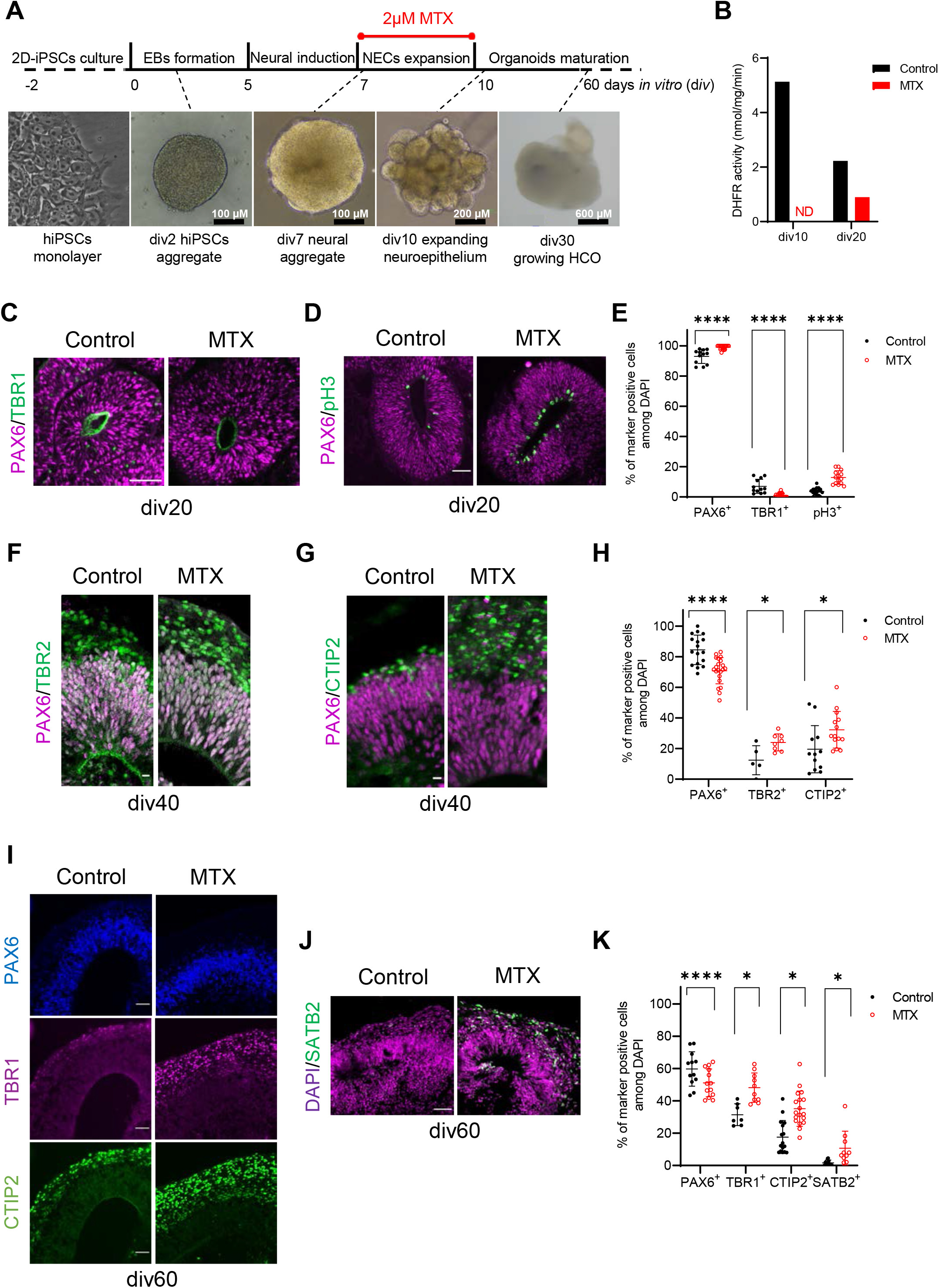
DHFR deficiency accelerate neurogenic program in HCO. (A) Timeline of the protocol used to generate HCO. The time window of MTX treatment is indicated. (B) DHFR metabolic activity in pooled DMSO (control) or MTX-treated HCO (n=8) at 2 different time points. (C-E) Representative images and quantification of control and MTX-treated HCO immunostained for pH3, PAX6 and TBR1 at div20. Scale bars, 50 μm. (F–H) Representative images and quantification of control and MTX-treated HCO immunostained for PAX6, TBR2 and CTIP2 at div40. Scale bars, 10 μm. (I-K) Representative images and quantification of control and MTX-treated HCO immunostained for PAX6, TBR1 CTIP2 and SATB2 at div60. Scale bars, 50 μm. Data are presented as mean ± SD (n = 7–12 organoids with 15–20 cortical-like structures analyzed with multiple Mann-Whitney test). ND, not detected.

### Engineering a *Dhfr* mutant mouse line

To investigate the role of DHFR on neurogenesis *in vivo* and the long-term consequences of its inhibition on neocortex development, we turned to the mouse as a model. Similar to Human, *Dhfr* mRNA is expressed in apical progenitors at early stages of mouse neocortex development when direct neurogenesis is prominent (Figure S2A). Using CRISPR/Cas9 engineering we generated a *Dhfr* mutant mouse line by deleting exon 5 of the *Dhfr* gene which harbors the catalytic activity (Figure 2A). Genotyping of E4.5 embryos revealed a wild type and a deleted allele indicating that genome editing was successful (Figure S2B) and RT-PCR and qRT-PCR analyses on WT and *Dhfr^+/Δ^* heterozygous E12.5 embryos showed that the deleted allele is transcribed (Figures S2C, D). *Dhfr^+/Δ^* heterozygous mutants were viable and fertile with normal body weight (Figure 2B). Genotyping at weaning age of pups born from heterozygous parents revealed that 31.98% (n=47) were WT and 68.02% (n=100) were *Dhfr^+/Δ^* heterozygous mutants. No homozygote mutants were collected past E4.5 suggesting a peri-implantation lethality. At E12.5, *Dhfr^+/Δ^* heterozygous embryos showed no gross morphological defects (Figure 2C). In tissue extracts from E12.5 embryos, DHFR protein level was decreased by half in *Dhfr^+/Δ^* samples compared to WT (Figure 2D). Decreased DHFR protein level correlated with a two-fold decrease in DHFR activity in *Dhfr^+/Δ^* samples compared to WT (Figure 2E). Altogether these data reveal that we generated a *bona fide* mouse model of DHFR deficiency.

**Figure 2.**
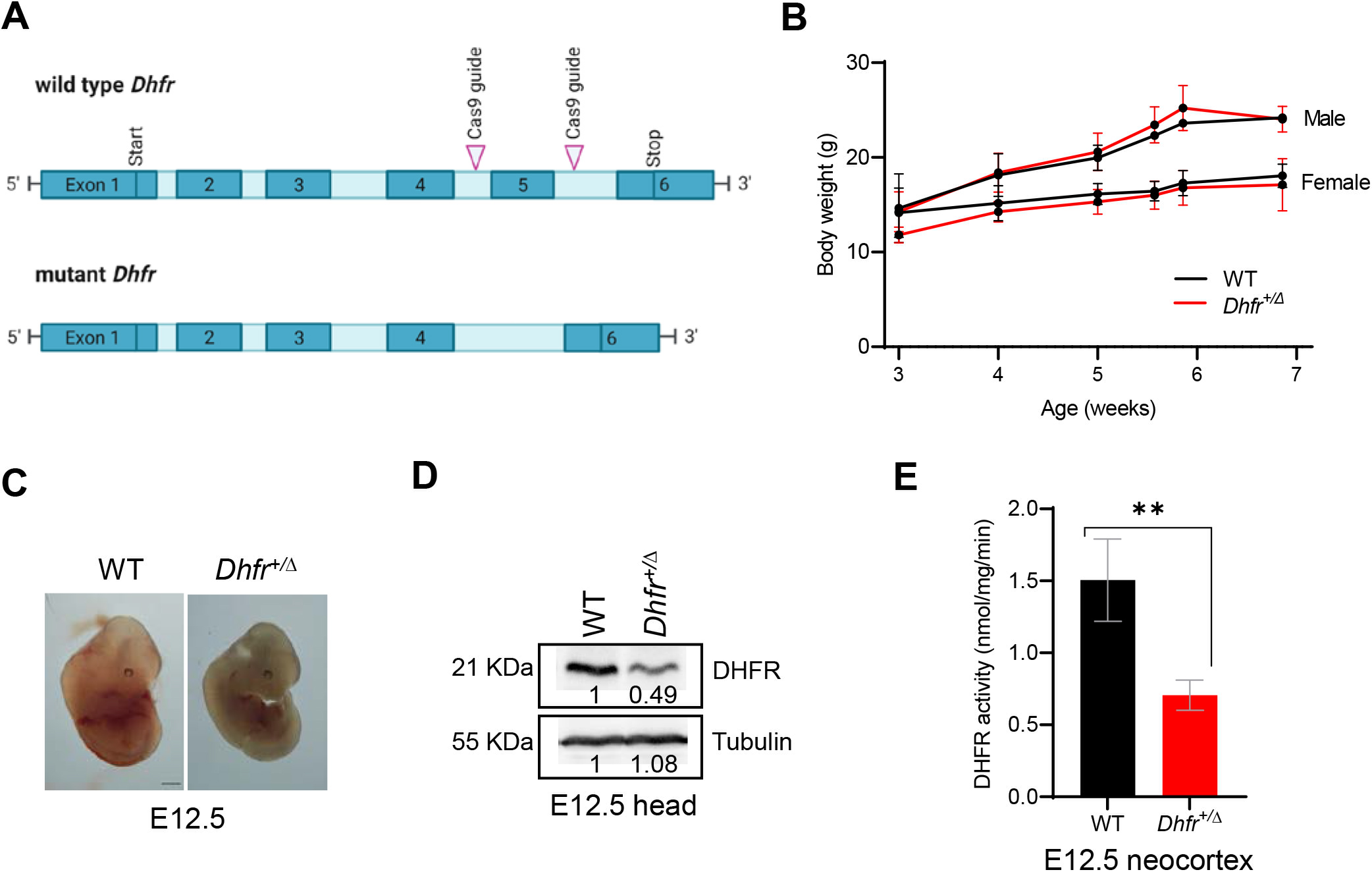
*Dhfr* heterozygous mutant mice exhibits reduced DHFR activity. (A) Schematic representation of the wild type *Dhfr* gene and the mutant allele obtained by deleting exon5 with CRISPR-Cas9 engineering. (B) Measurement of body weight of WT and *Dhfr^+/Δ^* pups from weaning to adult stage. (C) Representative images of WT and *Dhfr^+/Δ^* developing embryos at E12.5. (D) DHFR protein levels were analyzed by western blot analysis of WT and *Dhfr^+/Δ^* E12.5 head extracts (n=2). Tubulin was used as a loading control. Numbers indicate relative levels. (E) DHFR activity in WT and *Dhfr^+/Δ^* E12.5 neocortex (n=4). Data are presented as mean ± SD and statistical analysis was performed using unpaired t test or Mann-Whitney test.

### DHFR haploinsufficient embryos exhibit initial neocortex developmental delay

To gain insights into the role of DHFR in early neocortex development, we assessed progenitor and neuron populations in the *Dhfr^+/Δ^* heterozygous mutants at E12.5. This revealed that the number of PAX6^+^ apical progenitors was reduced in the lateral neocortex of *Dhfr^+/Δ^* embryos compared to WT embryos, with no change in TBR2^+^ basal progenitors (Figures 3A, D). In addition, we used TBR1, a marker of immature neurons (predominantly layer VI), CTIP2 (a marker of layer V neurons mostly produced by direct neurogenesis), and SATB2 (predominantly layer II-IV mostly produced by indirect neurogenesis) to quantify the production of projection neuron subtypes. In E12.5 *Dhfr^+/Δ^* embryos, the number of TBR1^+^ neurons was unchanged, the number of CTIP2^+^ neurons was significantly decreased in lateral neocortex and no SATB2^+^ neurons could be detected (Figures 3B, C, E). To understand the origin of the decrease in progenitors and CTIP2^+^ neurons, we examined progenitor proliferation and observed no significant differences in pH3-positive cells at the VZ of *Dhfr^+/Δ^* mice compared to WT (Figures 3F, G), suggesting unchanged proliferation rate of apical progenitors. In addition, immunostaining for the apoptotic marker cleaved Caspase-3 (CASP3) detected only a few condensed nuclei or CASP3-positve cells in WT and *Dhfr^+/Δ^* neocortex (Figures 3H, I), indicating that the decrease in progenitor and neuron populations is not due to cell death. Lastly, we quantified the mode of apical progenitor division since this could influence their number. Indeed, at early stages of development, most divisions of PAX6^+^ apical progenitors occur with a vertical cleavage plane, corresponding to amplifying divisions while divisions with horizontal cleavage plane corresponding to asymmetric neurogenic divisions are less frequent (Haydar et al., 2003). At E12.5, we observed that the proportion of divisions with a vertical plane (symmetric) is increased in *Dhfr^+/Δ^* neocortex compared to WT (Figure 3J) which does not fit with the observed smaller number of PAX6^+^ progenitors. Because it is known that the proportion of symmetric divisions decreases and the proportion of neurons increases as development proceeds, we thus hypothesized that all the phenotypes observed at E12.5 in *Dhfr^+/Δ^* mutants were in fact due to a global delay in neocortex development. To confirm this, we performed bulk RNA-sequencing at E12.5, comparing WT and *Dhfr^+/Δ^* embryonic neocortex. PCA analyses with the entire gene set showed that the samples could be partly clustered by genotypes (Figure S3A). Next, we performed a PCA biplot analysis using a subset of 258 genes specific for E12.5 neocortex (Di Bella et al., 2021) and observe an association of the majority of these genes with WT samples (Figure S3B) suggesting that the main difference in expression profiles between genotypes is related to genes defining this embryonic stage. Further bioinformatics analyses identified about 250 genes that are differentially expressed in *Dhfr^+/Δ^* embryonic neocortex (Figure S3C). We confirmed the molecular signature of an immature brain with decreased expression of *Ttr* and *Aldoc* by qRT-PCR (Figure S3D). Altogether, these results highlight the delayed progression of early neocortical development in *Dhfr^+/Δ^* embryos, which is reminiscent of the initial neurogenic delay observed in MTX-treated HCO.

**Figure 3.**
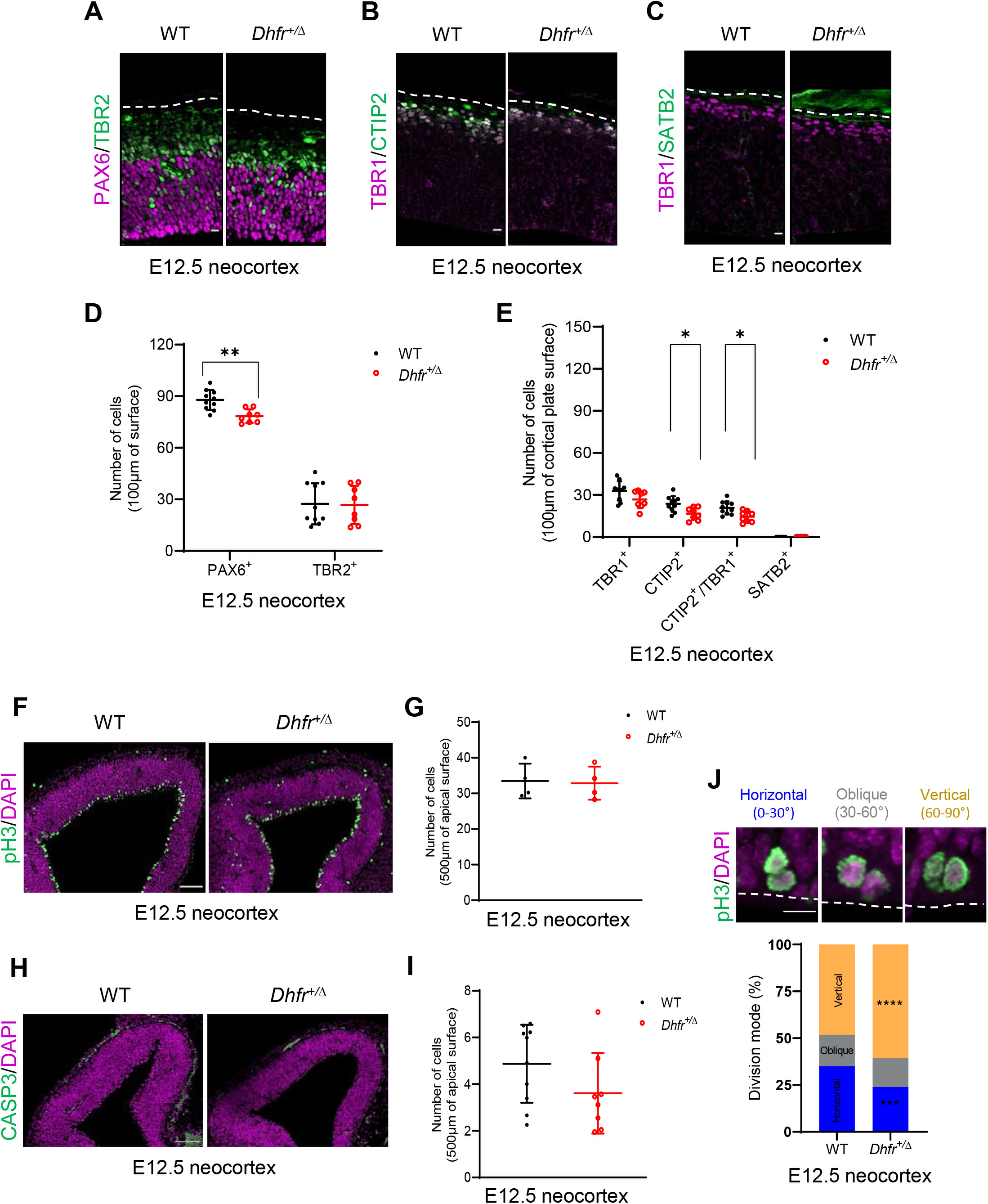
Initial developmental delay in *Dhfr* deficient embryos. (A-E) Representative images and quantification of neocortex coronal sections of E12.5 WT and *Dhfr^+/Δ^* embryos immunostained for PAX6, TBR2, TBR1, CTIP2 and SATB2. Scale bars, 10 μm. (F, G) Representative images and quantification of neocortex coronal sections of E12.5 WT and *Dhfr^+/Δ^* embryos immunostained for pH3. Scale bars, 100 μm. (H, I) Representative images and quantification of neocortex coronal sections of E12.5 WT and *Dhfr^+/Δ^* embryos immunostained for CASP3. Scale bars, 100 μm. (J) Top: Representative images of neocortex coronal sections of E12.5 WT and *Dhfr^+/Δ^* embryos immunostained for pH3. Images show examples of horizontal, oblique or vertical cleavage plane. Bottom: Quantification of cleavage plane orientation in the neocortex of WT and *Dhfr^+/Δ^* E12.5. Scale bars, 10 μm. Data are reported as mean ± SD and statistical analysis was performed using multiple unpaired t test or multiple Mann-Whitney test or two-way ANOVA test.

### DHFR activity is required to delay indirect neurogenesis in mouse neocortex

To investigate the consequences of reduced DHFR activity at later stages of mouse neocortex development, we quantified progenitor and neuron populations at E14.5 in WT and *Dhfr^+/Δ^* embryos. While the number of PAX6^+^ apical progenitors was unchanged, the number of TBR2^+^ basal progenitors was increased in E14.5 *Dhfr^+/Δ^* lateral neocortex (Figures 4A, D), suggesting that TBR2^+^ basal progenitors were produced at a higher rate from PAX6^+^ apical progenitors between E12.5 and E14.5. Concerning neurons, the number of TBR1^+^ neurons was unchanged in lateral neocortex, but the number of CTIP2^+^ neurons was decreased while the number of SATB2^+^ neurons was increased (Figures 4B, C, E), indicating a potential switch between direct neurogenesis (generating CTIP2^+^ neurons) and indirect neurogenesis (generating SATB2^+^ neurons from TBR2^+^ progenitors). Lastly, the number of pH3-positive cells within the SVZ was increased (Figures 4F, G), corresponding to the increased abundance of TBR2^+^ basal progenitors. At E16.5, the number of PAX6^+^ and TBR2^+^ progenitors are indistinguishable between WT and *Dhfr^+/Δ^* neocortex (Figures S4A, B), however, the imbalance in neuronal composition persists with fewer CTIP2^+^ and excess SATB2^+^ neurons (Figures S4C-F). No significant difference in neocortical thickness was observed between WT and *Dhfr^+/Δ^* heterozygous coronal sections (Figures S4G, H), indicating that general growth of the neocortex is not impaired in *Dhfr^+/Δ^* mutant embryos. Noticeably, this altered neuronal composition was still observed in *Dhfr^+/Δ^* somatosensory neocortex at postnatal P21 stage (Figures S4I, J) with a decreased number of CTIP2^+^ neurons and an increased number of SATB2^+^ neurons, revealing permanent changes in neuronal composition. These results indicate that DHFR activity is required to delay the onset of indirect neurogenesis in the mouse neocortex thus shaping the composition of neuronal subtypes present at adult stage.

**Figure 4.**
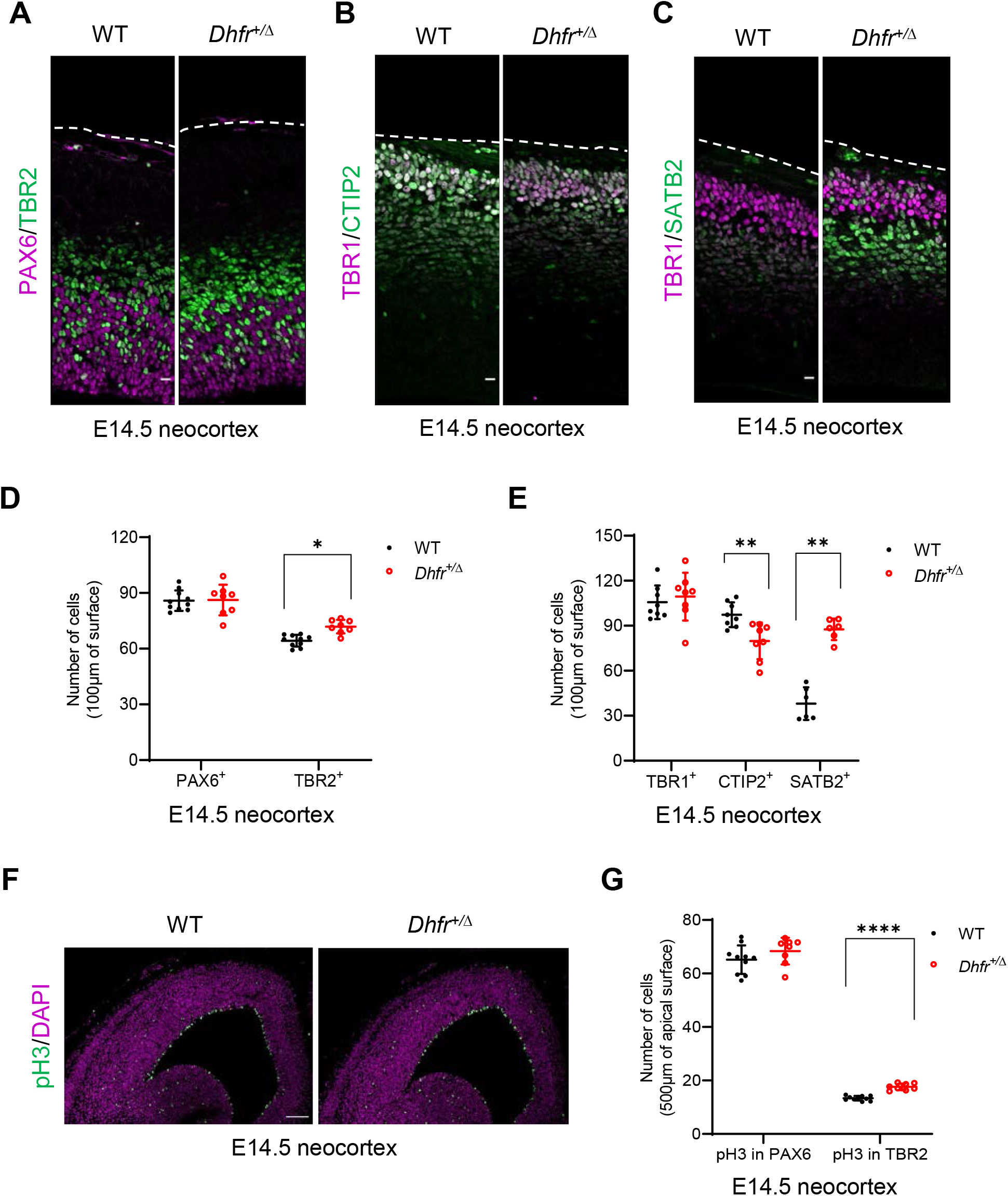
Overproduction of basal progenitors and late-born neurons in *Dhfr* deficient embryos. (A-E) Representative images and quantification of neocortex coronal sections of E14.5 WT and *Dhfr^+/Δ^* embryos immunostained for PAX6, TBR2, TBR1, CTIP2 and SATB2. Scale bars, 10 μm. (F, G) Representative images and quantification of neocortex coronal sections of E14.5 WT and *Dhfr^+/Δ^* embryos immunostained for pH3. Scale bars, 100 μm. Data are reported as mean ± SD and statistical analysis was performed using multiple unpaired t test or multiple Mann-Whitney test.

### DHFR deficiency modulates H3K4me3 marks

DHFR is a key enzyme in the One-carbon/folate metabolic pathway which has two distinct outputs: purine synthesis which is required for DNA synthesis and cell proliferation and synthesis of methyl groups which are required for methylation reactions (Figure 5A). Because we did not detect changes in progenitor proliferation that would be consistent with defective purine synthesis in Human or mouse neocortex, we hypothesized that DHFR deficiency affects neurogenesis by modulating the production of methyl groups and methylation reactions. To test this, we performed biochemical assays to measure levels of THF and of the universal methyl donor SAM and observed that both metabolites were significantly reduced in E12.5 *Dhfr^+/Δ^* head extracts compared to WT samples (Figures 5B, C). In a previous study we showed that H3K4me3 methylation marks were sensitive to DHFR inhibition in neural progenitors (Fawal et al., 2018). Investigation of this methylation mark by western blot analysis of mouse *Dhfr^+/Δ^* and Human neuroepithelial samples revealed a global loss of H3K4me3 marks in samples deficient for DHFR activity (Figure 5D). To test whether DHFR deficiency led to changes in H3K4me3 marks at specific genes, we performed H3K4me3 ChIP-sequencing on mouse neural progenitors treated for 72h with MTX *in vitro* and analyzed a set of neocortical layer-specific genes (Molyneaux et al., 2007). We noticed a reduction in H3K4me3 marks on a subset of deep layer-specific genes while the majority of upper layer-specific genes exhibited a slight increase in H3K4me3 marks (Figure 5E). These findings suggest that DHFR deficiency alters epigenetic landscapes in neural progenitors and this correlates with a switch in neuronal subtype identity generated. Importantly, these ChIP-Seq experiments were performed during a short time window on neural progenitors *in vitro,* leading to the conclusion that the observed changes in the level of H3K4me3 are not a consequence of developmental delay nor of spontaneous differentiation, they are a direct consequence of inhibiting DHFR activity in neural progenitors.

**Figure 5.**
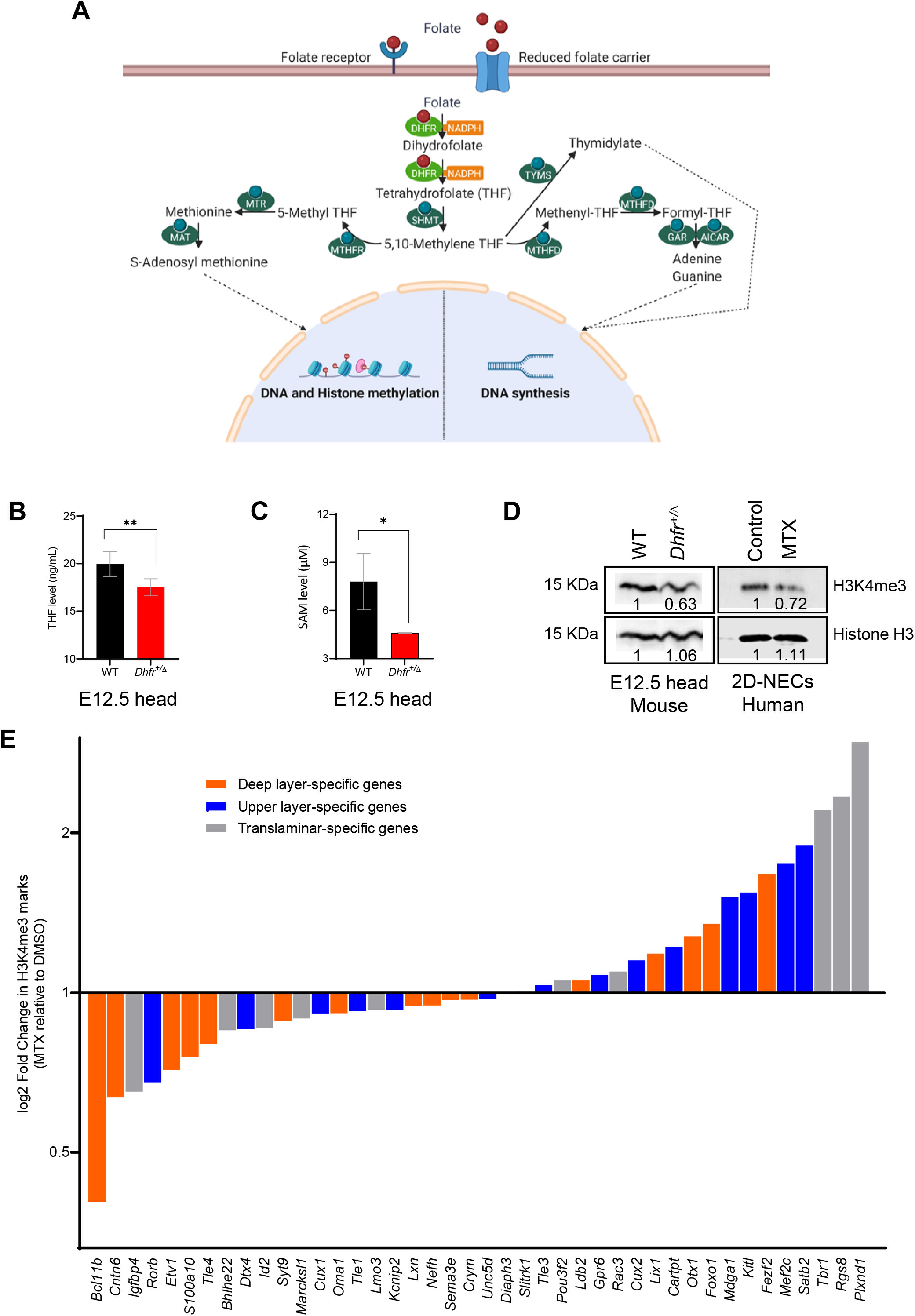
DHFR inhibition modifies H3K4 trimethylation. (A) Schematic representation of the 1C metabolic pathway and its outputs. (B, C) Levels of THF and SAM metabolites in WT and *Dhfr^+/Δ^* E12.5 head extracts (n=4-6 and n=3, respectively). (D) Left: H3K4me3 levels were analyzed by western blot analysis of WT and *Dhfr^+/Δ^* E12.5 head extracts (n=2). Right: H3K4me3 levels were analyzed by western blot analysis of 2D-NECs treated with DMSO (control) or with MTX. Histone H3 was used as a loading control. Numbers indicate relative levels. (E) Genome-wide H3K4me3 ChIP-sequencing data shows specific changes in methylation levels of neocortical layer-specific genes following 72 hours of MTX treatment in mouse neurosphere cultures. Data are reported as mean ± SD and statistical analysis was performed using multiple unpaired t test.

## DISCUSSION

Here we show that DHFR activity plays a role in controlling neurogenesis in the developing mammalian neocortex. First, we report the generation of *Dhfr^+/Δ^* haploinsufficient embryos which exhibit alteration in neurogenesis and neuronal subtype production. In the *Dhfr^+/Δ^* mutant mouse line, DHFR protein levels and DHFR activity are reduced by half in the embryonic neocortex revealing an absence of compensatory mechanisms from the wild type allele. Absence of compensatory mechanisms also applies to Humans, indeed, a two-fold reduction in DHFR protein levels and activity was reported in individuals carrying a heterozygous point mutation in *DHFR* (Banka et al., 2011; Cario et al., 2011). Although it is generally admitted that metabolic pathways are not sensitive to gene dosage due to buffering by other components of the network, this is not true for rate-limiting enzymes (Johnson et al., 2019). Despite the important role of DHFR in DNA replication, we observed no defect in proliferation and no increased cell death in the neocortex of *Dhfr^+/Δ^* heterozygous embryos, indicating that one copy of wild type *Dhfr* is sufficient to sustain purine synthesis. Conversely, we observed decreased levels of SAM and H3K4 trimethylation in *Dhfr^+/Δ^* embryonic neocortex indicating that DHFR activity is rate-limiting for the methionine cycle and methyl group synthesis in the neural tissue. In a previously described *Dhfr* mutant mouse line (*Ora*) generated by ENU mutagenesis, it was reported that heterozygous animals, which exhibited reduced DHFR activity, survived to adulthood with tissue-specific alterations in folate abundance and distribution, and perturbed stress erythropoiesis (Thoms et al., 2016). This data, together with our data, raises the possibility that *Dhfr* is an haploinsufficient gene and suggests that Human carriers of heterozygous mutations may have subtle alterations of the blood and brain compartments.

Recent single-cell RNA-sequencing data showed that *Dhfr* mRNA is highly expressed in neural progenitors at early stages of development and is down-regulated in apical progenitors at later stages, as well as in intermediate progenitors and differentiating neurons (Di Bella et al., 2021; Kanton et al., 2019; Telley et al., 2019). This dynamic expression pattern suggests that DHFR may play a specific function in apical progenitors. Analyses of neocortex development in *Dhfr^+/Δ^* haploinsufficient mouse embryos revealed an initial developmental delay followed by an acceleration of indirect neurogenesis culminating in the overproduction of late-born neurons at the expense of early-born neurons. Accelerated indirect neurogenesis accompanied by depletion of apical progenitors was also observed in HCO treated with MTX at an early stage of development indicating that these phenotypes are a direct consequence of DHFR inhibition in neural progenitors. These phenotypes are reminiscent of microcephalic features observed in patients harboring mutations in human *DHFR* (Banka et al., 2011) which could be due to premature differentiation leading to depletion of neural progenitor pools. Recent work studying the impact of folate-deficient diet on foetal corticogenesis in the mouse reported alterations in the production of neuronal subtypes that are similar to those observed in *Dhfr^+/Δ^* haploinsufficient embryos (Harlan De Crescenzo et al. 2021). However, the impact of folate-deficient diet on progenitors was different and was associated with increased cell death in the neocortex (Harlan De Crescenzo et al. 2021), suggesting that folate-deficient diet could have broad deleterious consequences on the pregnant dam that indirectly impact foetal corticogenesis.

It is well established that developmental transitions occurring during neocortex development involve chromatin modifications in neural progenitors (Albert et al., 2017; Hirabayashi and Gotoh, 2010). Here we show that inhibition of DHFR in mouse neural progenitors *in vitro* leads to discrete changes in H3K4me3 marks on neuronal specification genes. Indeed, following MTX treatment, a number of genes expressed in early-born neurons have less H3K4me3 while some genes expressed in late-born neurons have more, which is consistent with the neuronal imbalance observed *in vivo* in *Dhfr^+/Δ^* embryos. These results suggest that variations in DHFR activity influence the epigenetic landscape in apical progenitors and that these epigenetic modifications set the stage for production of the different neuronal subtypes. Indeed, recent single-cell analyses of chromatin and gene-regulatory dynamics in the developing brain suggested that progenitors entering the cell cycle may be epigenetically primed toward future cell states (Trevino et al., 2021).

Our study identified a conserved function for DHFR in preventing indirect neurogenesis. Indeed, we observed an increased production of TBR2^+^ intermediate progenitors following DHFR inhibition both in mouse and Human contexts. While it is admitted that apical progenitors generate neurons both directly and indirectly, the molecular mechanisms governing the switch from one to the other neurogenic process are not well characterized. In the mouse it was estimated that 5% of apical progenitors produce neurons directly at E12.5 and that the switch between direct and indirect neurogenesis involves Robo and Dll1 signaling (Cárdenas et al., 2018). In the future it would be interesting to assess the link between DHFR and these signaling pathways. One defining feature of indirect neurogenesis is that it is a hallmark of mammalian neocortex evolution. Indeed, direct neurogenesis with limited neuronal production dominates avian and reptilian paleocortex while indirect neurogenesis is predominant in mouse neocortex (Cárdenas et al., 2018). In the Human neocortex indirect neurogenesis is vastly expanded and accounts for the massive increase in neuron numbers (Hansen et al., 2010; Lewitus et al., 2014; Namba and Huttner, 2017). Whether this evolutionary innovation was driven, at least in part, by evolutionary changes in 1C metabolism is an interesting matter for speculation.

## MATERIALS AND METHODS

All materials and methods are described in Supplemental Methods.

## ACKNOWLEDGEMENTS

We would like to thank Alain Vincent and Eric Agius for critical reading of the manuscript. We acknowledge the help and contribution of the ANEXPLO mouse facility and the TRI imaging platform at the CBI. We also thank the CBI-bigA facility engineers for their help and feedback during the raw data processing, the statistical analysis and the data visualization of sequencing data. This work was funded by Fondation pour la Recherche Médicale (DEQ20180339174) and Agence Française pour la Recherche (ANR-21-CE16-0024-01). Sequencing for the ChIP-Seq analysis was performed by the GenomEast platform, a member of the ‘France Génomique’ consortium (ANR-10-INBS-0009). Both CNRS and Université de Toulouse provided core funding. Sulov SAHA is the recipient of a 3-year PhD scholarship from the French Ministry of Research.

## AUTHOR CONTRIBUTIONS

SS performed experiments, analyzed and interpreted data, wrote a draft and edited the manuscript; TJ performed experiments, analyzed and interpreted data, edited the manuscript; DO performed bioinformatics analyses, interpreted data and edited the manuscript; CA performed experiments; TY performed bioinformatics analyses; MAF performed experiments, analyzed and interpreted data, edited the manuscript; AD conceptualized and managed the study, analyzed and interpreted data, wrote the manuscript.

## CONFLICT OF INTEREST

The authors declare no conflict of interest.

## Notes

### Competing Interest Statement

The authors have declared no competing interest.

