## Supplemental figures for "DHFR metabolic activity controls neurogenic transitions in the developing Human and mouse neocortex"

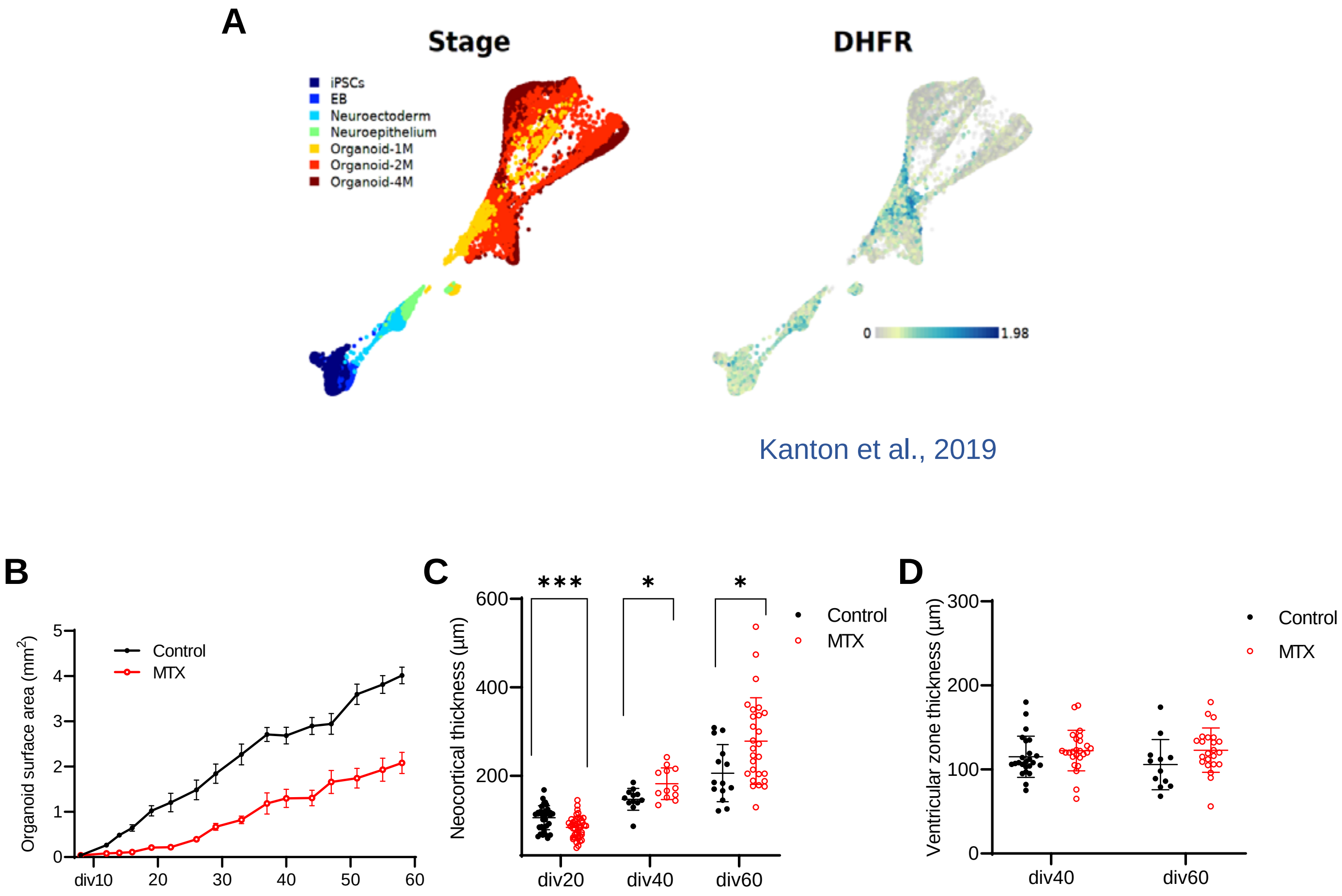


**Figure S1. *DHFR* expression and inhibition in HCO**

(A) *DHFR* mRNA expression over the course of HCO development and maturation (data collected from Kanton et al., 2019).

(B) Quantification of the size of control and MTX-treated HCO over time.

(C, D) Measurement of the relative thickness of the neocortex and ventricular zone in control and MTX-treated HCO.

Data are presented as mean ± SEM or mean ± SD (n = 7–12 organoids with 15–20 cortical-like structures analyzed with multiple Mann-Whitney test).

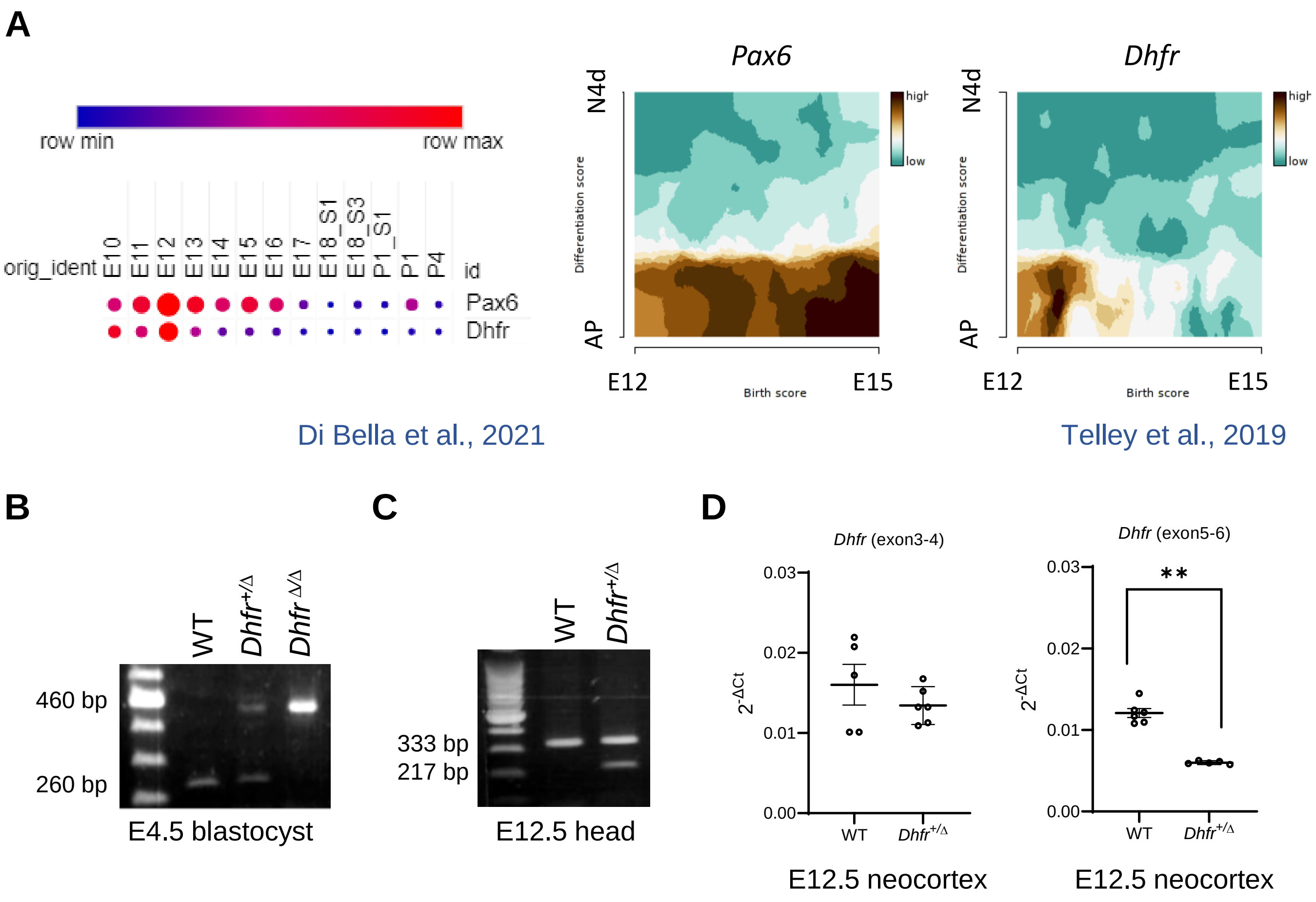


**Figure S2. *Dhfr* expression in wild type and mutant mice**

(A) *Dhfr* mRNA expression in the developing mouse neocortex (data collected from Di Bella et al., 2021; Telley et al., 2019). Abbreviations: AP, apical progenitors; E, embryonic day; N4d, 4-day-old neurons; P, postnatal day.

(B) Representative genotyping results of WT (*Dhfr*$^{+/+}$), *Dhfr*$^{+/\Delta}$ and *Dhfr*$^{\Delta/\Delta}$ E4.5 embryos. The 260 bp band corresponds to the wild type allele and the 460 bp band corresponds to the edited allele.

(C) RT-PCR on mRNA extracted from WT and *Dhfr*$^{+/\Delta}$ E12.5 embryos was used to detect expression of the wild type and edited alleles of *Dhfr*.

(D) qRT-PCR analysis of *Dhfr* expression. Two sets of primers were used to amplify either exon3-4 (not sensitive to the excision, amplification from the two alleles) or exon5-6 (sensitive to the excision, amplification from the wild type allele only).

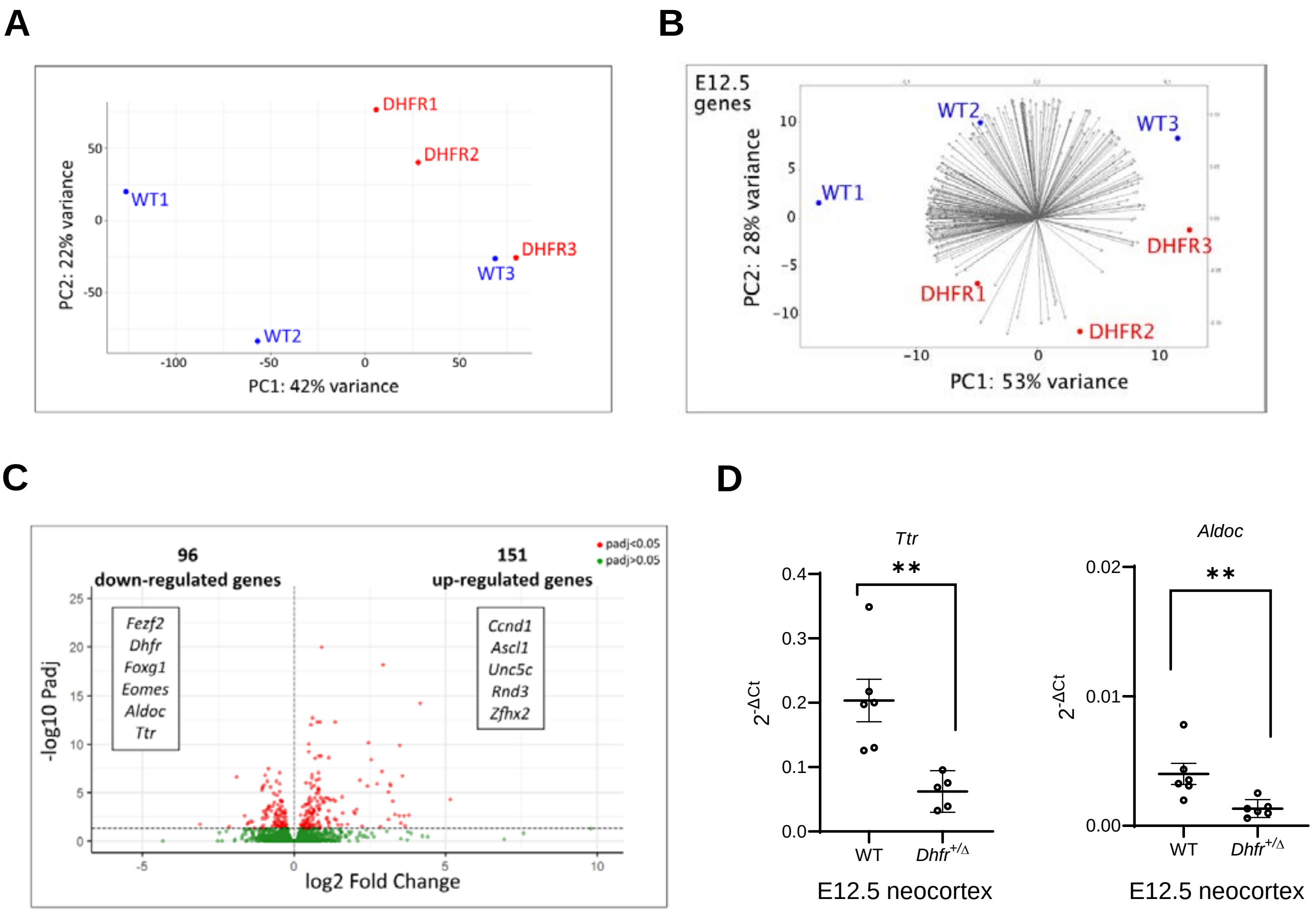


**Figure S3. Bulk RNA-sequencing reveals developmental delay in E12.5 *Dhfr* deficient embryos**

(A) PCA plot showing the distribution of WT *(Dhfr$^{+/+}$)* and DHFR *(Dhfr$^{+/\Delta}$)* embryos based on whole genome expression profiles.

(B) PCA biplot showing the comparison of WT *(Dhfr$^{+/+}$)* and DHFR *(Dhfr$^{+/\Delta}$)* embryos based on a subset of 258 genes specific for E12.5 neocortex (from Di Bella et al., 2021).

(C) Numbers of genes significantly up-regulated and down-regulated in *Dhfr$^{+/\Delta}$* are indicated in red dots.

(D) qRT-PCR analysis of selected genes.

### Figure S4. *Dhfr* deficiency irreversibly alters mouse neocortical composition

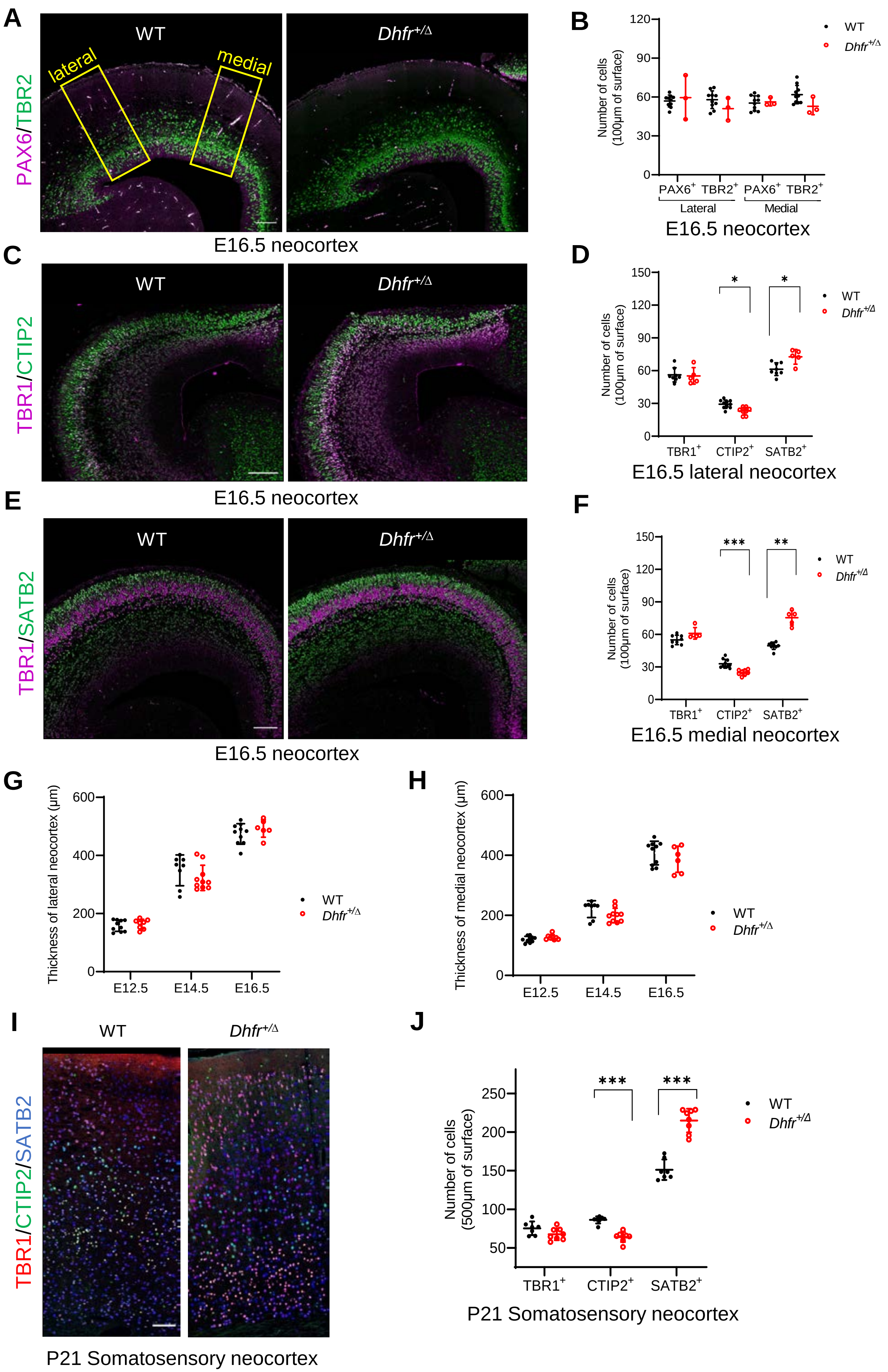

**Figure S4. *Dhfr* deficiency irreversibly alters mouse neocortical composition**

(A-F) Representative images and quantification of neocortex coronal sections of E16.5 WT and *Dhfr*$^{+/\Delta}$ embryos immunostained for PAX6, TBR2, TBR1, CTIP2 and SATB2. Regions of interest corresponding to medial and lateral neocortex (yellow box) were quantified. Scale bars, 100 μm.

(G, H) Measurement of relative neocortical thickness at medial and lateral positions.

(I, J) Representative images and quantification of coronal sections of somatosensory cortex of P21 WT and *Dhfr*$^{+/\Delta}$ pups immunostained for TBR1, CTIP2 and SATB2. Scale bars, 100 μm.

Data are reported as mean ± SD and statistical analysis was performed using multiple unpaired t test.
