## Supplemental Methods for "DHFR metabolic activity controls neurogenic transitions in the developing Human and mouse neocortex"

### Material and Methods

#### RESOURCE AVAILABILITY

##### CONTACT FOR REAGENT AND RESOURCE SHARING

##### MATERIALS AVAILABILITY

The mouse line generated in this study is available upon request.

##### DATA AVAILABILITY

RNA-Seq and ChIP-Seq data have been deposited at GEO and are publicly available as of publication. Accession numbers are listed in the key resources table. Original western blot and microscopy data will be shared by the lead contact upon request.

#### EXPERIMENTAL MODEL AND SUBJECT DETAILS

##### ANIMALS

Wild type and *Dhfr*$^{+/\Delta}$ mice were kept in a 129S4/C57Bl6J mixed background. Mice were maintained on a standard 12 hours day:night cycle and given ad libitum access to food and water. Mice were housed in ventilated cages, 3-5 animals per cage, except males used for mating (1 per cage). Timed pregnancies were obtained by detecting vaginal plugs. Embryos at stages from E12.5 to E16.5 and pups at P21 were analyzed. The gender of embryos was not tested, littermates of each genotype were randomly assigned to experimental groups. All experimental procedures were pre-approved and carried out in compliance with the guidelines provided by the national Animal Care and Ethics Committee (APAFIS#1289-2015110609133558 v5) following Directive 2010/63/EU.

##### PLURIPOTENT STEM CELLS

Human iPSCs were obtained from Leiden University Medical Center (LUMC0004iCTRL). Cells were amplified with mTesR+ medium (Cat#100-0276, STEMCELL Technologies) on h-ES qualified Matrigel (Cat#354277, Corning) coating and stored frozen with CryoStor CS10 (Cat#07930, STEMCELL Technologies) as a ready to work bank in liquid nitrogen.

##### PRIMARY MOUSE NEURAL PROGENITOR CELLS

Neocortical tissue from 4-6 E13.5 embryos were dissociated mechanically in PBS. The single-cell suspension was collected, rinsed with DMEM/F12 (Cat#1130-032, Invitrogen), and cultured with growing medium (DMEM/F12 medium containing 0.6% Glucose (Cat#UG3050, Euromedex), 5 mM HEPES (Cat#H3375, Sigma-Aldrich), 1 mM Putrescine (Cat#P5780, Sigma-Aldrich), 5 ng/ml FGF-2 (Cat#F029, Sigma-Aldrich), 20 ng/ml EGF (Cat#E9644, Sigma-Aldrich), 10 ng/ml Insulin-transferrin-sodium selenite supplement (Cat#I1844-1VL, Sigma-Aldrich) and 2% B27 supplement (Cat#17504-044, Invitrogen) in a 5% $CO_2$ incubator at 37°C. Neural progenitor cells were cultured using Nunc cell culture flasks (Cat#156340, Thermoscientific) as free-floating neurospheres for 72 hours.

#### METHOD DETAILS

##### Generation and maintenance of HCO

For every single batch of cerebral organoids one vial of same passage iPSCs was defrosted 3-4 days and cells were allowed to reach 70-80% confluence with daily changing medium in mTesR+. HCO were generated using STEMdiff cerebral organoid kit (Cat#8570, STEMCELL Technologies) and according to manufacturer recommendations. Briefly, iPSCs were dissociated on day 0 as single cells using GCDR solution (Cat#07174, STEMCELL Technologies) and 9000 cells were aggregated as Embryoid Bodies (EBs) in EB formation medium containing 10 µM ROCK-inhibitor (Y-27632, Cat#7302, STEMCELL Technologies) in U bottom 96-well plates (Cat#650970, Greiner) for 5 days when EBs are large enough (typically 400-600 µm) to be transferred in ultra-low attachment 48-well plates (Cat#150787, ThermoFisher) with neural induction medium for 2 days. At day *in vitro* 7 (div7) neuroepithelial aggregates where embedded in 15 µl of h-ES qualified Matrigel (Cat#354277, Corning) droplets and neuroepithelium buds expanded to div10 in expansion medium when early HCO are transferred in non-adherent 6-well plates (Cat#158476, ThermoFisher) under orbital agitation (Cat#A073545, Grosseron, set at 73 rpm) in maturation medium. HCO maturation was prolonged to div60 as final time point, with medium replacement every 3 days, orbital shaking and the STEMdiff cerebral organoid maturation kit medium (Cat#8571, STEMCELL Technologies) in a 5% $CO_2$ incubator (Forma Scientific) at 37°C. For acute inhibition of DHFR activity in HCO, neuroepithelial aggregates were treated at div7 with 2 µM MTX (Cat#M8407, Sigma-Aldrich) freshly diluted in expansion medium, for 3 days. At div10 medium was replaced as for control conditions with MTX-free maturation medium.

HCO were collected and fixed in 15 ml conical tubes with 4% Paraformaldehyde (Cat#15710, Fisher scientific) overnight at 4°C and transferred to 30% Sucrose (Cat#S0389, Sigma-Aldrich) solution freshly made in double-distilled water at 4°C. After sinking (1-3 days), HCO were embedded in OCT (Cat#4583, Sakura) in plastic mold (5-10 organoids per condition depending on size). They were stored at -80°C until serial 15 µm thickness sectioning was performed using Cryostat (Cat#CM1950, Leica Biosystems) and serial sections were collected on SuperFrost Plus slides (Cat#J1800AMNT, Thermoscientific). For immunostaining, sections were allowed to reach room temperature before incubate in blocking solution (1% BSA (Cat#04100811C, Euromedex), 1% FBS (Cat#10500064, ThermoFisher), 0.5% Tween 20 (Cat#2001-A, Euromedex) and PBS (Cat#D1408, Sigma-Aldrich)) for 30 minutes at room temperature. Then sections were covered with sufficient volume of blocking solution containing adapted dilutions of primary antibodies mix (Table S2) for an overnight at 4°C in wet atmosphere. After 3 washes for 5 minutes in PBS, sections were covered with sufficient volume of blocking solution containing adapted dilutions of secondary antibodies (Table S2) and DAPI (Cat#62248, ThermoFisher) mix for 2 hours at room temperature, washed again 3 times in PBS and mounted with coverslips (Cat#17234914, Menzel-Gläser) and mounting medium (4.8% wt/vol Mowiol (Cat#81381, Sigma-Aldrich) and 12% wt/vol Glycerol (Cat#G9012, Sigma-Aldrich) in 50 mM Tris [pH 8.5] (Cat#26-128-3094-B, Euromedex)).

##### 2D-neuroepithelial cells culture

2D-neuroepithelial cells (2D-NECs) were derived from human-iPSCs using the STEMdiff SMADi neural induction kit (Cat#08581, STEMCELL Technologies) and according to manufacturer recommendations for the monolayer culture protocol. Briefly, iPSCs initially grown in mTESR+ on coating Matrigel were subjected to neural induction for 3 consecutive passages using Accutase (Cat#07920, STEMCELL Technologies) and 21 days in total with STEMdiff SMADi neural induction medium. Then they were grown and amplified for 2 additional passages, using Accutase in STEMdiff neural progenitor medium (Cat#05833, STEMCELL Technologies) on Matrigel and stored frozen in liquid nitrogen using STEMdiff neural progenitor freezing medium (Cat#05838, STEMCELL Technologies). For inhibition of DHFR activity in 2D-neuroepithelial cells, 50% confluent epithelial cells were treated for 3 days with 2 µM MTX diluted in growing medium and cells processed for analysis immediately after.

##### Generating CRISPR/Cas9-mediated *Dhfr* deletion in mice

Using the publicly available design program CRISPOR (http://crispor.tefor.net), we designed 2 CRISPR-Cas9 crRNA targeting intronic sequences flanking *Dhfr* exon5: 5′-GGATTAGCAACTAGGTAATA-3′ and 5′-TCCAAGCTGGCCCCATGTTT-3′. Each crRNA was complexed with tracrRNA to generate gRNA complexes, which were used at a 1/1 molar ratio to form the ribonucleoprotein complex with Cas9 protein (all tools from Integrated DNA Technologies). Cas9-ribonucleoprotein complex was microinjected in 129S4/C57Bl6J mixed background 1-cell embryos. Injected embryos were re-implanted in pseudo-pregnant dams. Progeny were screened by DNA sequencing (primer Fw: 5′-GTGATTCCATAGCCTCTTCCAG-3′, Rev: 5′-TAGCACACATGCTCCTTTCC -3′). A heterozygote founder mouse was identified with a 844 bp deletion corresponding to *Dhfr* exon5. This founder mouse was crossed with 129S4/C57Bl6J mice to establish the *Dhfr*$^{+/\Delta}$ mouse line. We tested 2 off-targets site for each CrRNA guide in this mouse line. We determined the most predictive off-target site with CRISPOR program and designed primers to amplify the corresponding regions. By sequencing, we eliminated the likelihood of indel formation in the 4 off-target sites analyzed. All genome editing experimental procedures were pre-approved and carried out in compliance with the guidelines provided by the national Animal Care and Ethics Committee (APAFIS #11931 2017102511049339) following Directive 2010/63/EU.

##### Genotyping

Genomic DNA was isolated from mouse tail, and genotyping was performed using 2x PCR Taq MasterMix (Cat#G013-dye, abm) with primers listed in Table S1. The PCR amplification was performed using Veriti thermal cycler (Cat#4452300, ThermoFisher) followed by separation of DNA fragments on agarose gel (1.2%). Ethidium bromide (Cat#EU0070, Euromedex) was used to make DNA bands visible under UV light using a gel documentation system (Cat#InGenius, Syngene).

##### Histochemistry

Collected developing mouse brains were fixed with 4% Paraformaldehyde overnight at 4°C followed by paraffin embedding with Spin tissue processor (Cat#STP120, Myr) for coronal sections on Microtome (Cat#HM355, Microm) or agarose (5% low-melting) embedding for coronal sections on Vibratome (Cat#VT1000S, Leica biosystems). Paraffin-embedded sections were deparaffined, rehydrated, followed by antigen retrieval involving steam heating (Cat# HD9104/01,

Phillips) for 45 minutes. Coronal sections were permeabilized and blocked with PBTA solution (1-3% BSA, 1-3% FBS, 0.5-1% Tween 20 and PBS) for 2 hours at room temperature. Primary antibodies (Table S2) diluted in PBTA solution; secondary antibodies (Table S2) and DAPI diluted in PBT solution (0.5-1% Tween 20 and PBS) were incubated overnight at 4°C and 2-3 hours at room temperature, respectively. Mounted sections on SuperFrost Plus slides were secured with coverslips and mounting medium.

**RNA-sequencing**

Brains from E12.5 WT and *Dhfr*$^{+/\Delta}$ embryos were collected and the neocortex was dissected. The tissue was mechanically dissociated to form a cell pellet. RNA was isolated from the cell pellet using Trizol reagent (Cat#15596026, Invitrogen) according to manufacturer instructions and eluted into 35 µl Nuclease-free water (Cat#W4502, Sigma-Aldrich). Eluted RNAs were then stored at -80°C. RNA quality and integrity (RIN>7) were verified by electrophoresis using 2100 Bioanalyzer system (Cat#G2939BA, Agilent). The whole procedure going from the dissociation to RNA preparation was performed on 6 animals (3 WT and 3 *Dhfr*$^{+/\Delta}$) obtained from 2 different litters.

RNA libraries were prepared using NEBNext ultra II reagent (Cat#E7770S, New England Biolabs) and RNA sequencing on neocortical samples were performed by IntegraGen (https://integragen.com) using an Illumina NovaSeq 6000 S2 sequencing system (paired-end sequencing, 100 bp reads, 35M reads/sample). The quality of each raw sequencing file (fastq format) was verified with FastQC analysis (https://www.bioinformatics.babraham.ac.uk/projects/fastqc/). All files were aligned to the reference mouse genome (Ensembl *Mus musculus* GRCm39; gtf(mm10)) using RNA STAR (Version Galaxy Tool: v2.7.2b; Dobin et al., 2012). Read count per sample was computed using HT-seq count (Version Galaxy Tool: v0.6.1 galaxy3; (Anders et al., 2015)). The raw count table was cleaned, genes having a read count summed higher or equal to 120 were kept for further analysis.

Differential analysis was applied per genotype (WT vs *Dhfr*$^{+/\Delta}$) by taking into consideration of litter effects using DESeq2 (v1.32.0; (Love et al., 2014)), available as an R package in Bioconductor (www.biocondutor.org). The raw read count were normalized using RLE methods generating and the log2 Fold Change (log2FC) values were computed. We used principal component analysis (PCA) to cluster samples based on their expression levels. Data representation of the scaled PCA was generated using the ggplot2 package (https://ggplot2.tidyverse.org).

**ChIP-sequencing**

E13.5 neocortical tissues were dissociated mechanically in PBS. The single-cell suspension was collected, rinsed with DMEM/F12 (Cat#1130-032, Invitrogen), and cultured with growing medium (DMEM/F12 medium containing 0.6% Glucose (Cat#UG3050, Euromedex), 5 mM HEPES (Cat#H3375, Sigma-Aldrich), 1 mM Putrescine (Cat#P5780, Sigma-Aldrich), 5 ng/ml FGF-2 (Cat#F029, Sigma-Aldrich), 20 ng/ml EGF (Cat#E9644, Sigma-Aldrich), 10 ng/ml Insulin-transferrin-sodium selenite supplement (Cat#I1844-1VL, Sigma-Aldrich) and 2% B27 supplement (Cat#17504-044, Invitrogen) in a 5% $CO_2$ incubator at 37°C. Neural progenitor cells were cultured using Nunc cell culture flasks (Cat#156340, Thermoscientific) as free-floating neurospheres for 72 hours. 2 µM MTX were diluted in growing medium for inhibiting DHFR activity. Cells were fixed with 1% Formaldehyde (Cat#15710, Fisher Scientific) for 15 minutes at room temperature, with occasional swirling. Glycine (Cat#G7126, Sigma-Aldrich) was added to a final concentration of 0.125 M and the incubation was continued for an additional 5 minutes. Cells were collected and washed with ice-cold PBS and resuspended in cell lysis buffer (5 mM Pipes [pH 8] (Cat#P6757, Sigma-Aldrich), 85 mM KCl (Cat#P017-A, Euromedex), 0.5% NP-40 (Cat#N6507, Sigma-Aldrich), Protease inhibitors (Cat#11836170001, Roche). Cells were centrifuged (Cat#ST40R, Eppendorf) at 4,000 rpm for 10 minutes at 4°C. Pellets were then resuspended in Nuclei lysis buffer (50 mM Tris-HCl [pH 8] (Cat#EU0011, Euromedex), 10 mM EDTA (Cat#EU0007, Euromedex), 1% SDS (Cat#EU0660, Euromedex)). The cells were disrupted by sonication (30 cycles, 30 seconds on, 60 seconds off) with Biorupter sonicator (Cat#B01020001, Diagenode). The chromatin solution was clarified by centrifugation at 15,000 g at 4°C for 10 minutes. The average DNA fragment size was 250 pb. The chromatin solution was diluted with IP dilution buffer (16.7 mM Tris-HCl [pH 8], 1.2 mM EDTA, 1.1 mM Triton X-100 (Cat#T9284, Sigma), 0.01% SDS and 167 mM NaCl (Cat#1112-A, Euromedex) and precleared with pre-blocked beads (protein G Sepharose (Cat#3296, Sigma-Aldrich)/protein A Agarose (Cat#6526, BioVision) (50/50) overnight with PBS/0.5% BSA, 10 mg/ml yeast tRNA (Cat#AM7119, ThermoFisher) beads for 1 hour at 4°C. The precleared diluted chromatin sample was incubated with 3 mg of anti-H3K4me3 (Cat#ab8580, Abcam) overnight at 4°C. Preblocked beads were added for an additional 4 hours. The beads were washed 2 times with the dialysis buffer (2 mM EDTA, 50 mM Tris-HCl [pH 8] and 0.2% N-lauroylsarcosine (Cat#L5777, Sigma-Aldrich), 4 times with wash buffer (100 mM Tris-HCl [pH 8], 500 mM LiCl (Cat#L4408, Sigma-Aldrich), 1% NP-40 and 1% Sodium deoxycholate (Cat#D6750, Sigma-Aldrich) and 2 times with TE buffer (10 mM Tris-HCl [pH 8] and 1 mM EDTA). The immunoprecipitated material was eluted from the beads by heating for 15 minutes at 65°C in 1% SDS, 50 mM $NaHCO_3$ (Cat#71631, Sigma-Aldrich). 0.2 M NaCl and 10 mg/ml RNase A (Cat#EN0531, ThermoFisher) were added before incubating for 2 hours at 64°C to reverse the crosslinks. Samples were then incubated with 1.5 mg/ml Proteinase K (Cat#P6556, Sigma-Aldrich), 40 mM Tris-HCl [pH 8], 10 mM EDTA at 45°C for 1 hour. The DNA samples were then extracted with Phenol chloroform isoamyl alcohol (Cat#0038.2, Carl Roth) followed by Ethanol (Cat#20821.330, VWR chemicals) precipitation in the presence of Glycogen (Cat#R0561, ThermoFisher), and resuspended in Nuclease-free water.

GenomEast (http://genomeast.igbmc.fr) performed DNA quantification with Qubit (Invitrogen) and processed 1.5-10 ng of double-stranded purified DNA to generate libraries using MicroPlex library preparation kit v2 (Cat#C05010012, Diagenode). Amplified libraries were purified with Agencourt AMPure XP beads (Cat#A63880, Beckman Coulter) and sequenced on an Illumina Hiseq 4000 sequencer (single-end sequencing, read length 50, minimum 45M reads/sample). Sequence reads were mapped to the *Mus musculus* genome assembly mm10 using Bowtie (Langmead et al., 2009). Peak calling was done by MACS v2.1.1 (Zhang et al., 2008; Feng et al., 2012). Peaks were annotated to the Ensembl release 94 using HOMER software (http://homer.ucsd.edu/homer/ngs/annotation.html). Peaks from different conditions and replicates were merged to form a consensus peak set. The read coverage for each sample was calculated with multicov function from bedtools v2.26.0 (Quinlan and Hall, 2010). Differential analysis was applied per condition (DMSO vs MTX) using DESeq2 (v1.20.0; Love et al., 2014) to calculate log2 Fold Change of H3K4me3 ChIP-seq read counts between MTX condition versus

DMSO condition. Data representation of the differential binding analysis corresponds to a strong change in peak heights at the promoter or gene body.

##### RT-PCR and qRT-PCR

RNA was extracted from E12.5 head (for RT-PCR) or E12.5 neocortex (for qRT-PCR) using Trizol reagent according to the manufacturer's instructions. 1 µg RNA was used for reverse transcription. Genomic DNA was degraded with 1 µl DNase (Cat#M6101, Promega) for 20 minutes at 37°C in 20 µl Nuclease-free water, and the reaction was stopped by adding 1 µl stop solution under heat inactivation at 65°C for 10 minutes. 2 µl of 10 mM dNTPs (Cat#U1511, Promega) and 2 µl of 100 mM OligodTs (Integrated DNA Technologies) were added for 5 minutes at 65°C, then 8 µl of 5x buffer, 2 µl RNasin (Cat#N2511, Roche), and 4 µl of 100 mM DTT (Cat#P1171, Promega) were added for 2 minutes at 42°C. The mix was divided into equal volumes in a reverse-transcriptase–negative control tube with addition of 1 µl Nuclease-free water and in a reverse-transcriptase–positive tube with 1 µl Superscript enzyme and placed at 42°C for 1 hour. The reaction was stopped at 70°C for 15 minutes, and cDNAs were synthesized.

For RT-PCR, 1 µl diluted cDNA (10-fold) was mixed with 2x PCR Taq MasterMix (Cat#G013-dye, abm) containing 1 µM of each primer listed in Table S1, and PCR amplification was performed using Veriti thermal cycler (Cat#4452300, ThermoFisher) followed by separation of DNA fragments on agarose gel (1.2%). Ethidium bromide (Cat#EU0070, Euromedex) was used to make DNA bands visible under UV light using gel documentation system. A no template reaction was used as a negative control.

For quantitative PCR, cDNAs were diluted (10-, 100-, and 1,000-fold) and processed in triplicate for each dilution. 10 µl diluted cDNA was mixed with 10 µl premix Evagreen (Cat#1725204, Bio-Rad) containing 1 µM of each primer, and the PCR program was run for 40 cycles on CFX96 Real-Time PCR system (Cat#185-5096, Bio-Rad). mRNA relative expression levels were calculated using the $2^{-\Delta Cts}$ method with primers listed in Table S1.

##### Western blot

For protein extraction, mouse and Human tissues were obtained by adding lysis buffer (150 mM NaCl, 50 mM Tris-HCl [pH 7.4], 0.5 mM EDTA, 2 mM $Na_3VO_4$ (Cat#S6508, Sigma-Aldrich), 1% NP-40, 0.5 mM EGTA and 0.1 mM PMSF (Cat#78830, Sigma-Aldrich) supplemented with protease inhibitors for 1 hour on ice. Protein lysates were vortexed, ultrasonicated for 5 cycles (Cat#11862075, Bioblock), and centrifuged (Cat#5415R, Eppendorf) at 1,500 rpm for 10 minutes. Protein concentration in the supernatant was determined using DC protein assay (Cat#5000112, Bio-Rad) based on colorimetric analysis with Spectrophotometer (Cat#720001, Jenway). Lysates were then denatured by boiling in 4x loading buffer (100 mM Tris-HCL [pH 6.8], 8% SDS, 40% Glycerol, 4% β-mercaptoethanol (Cat#63689, Sigma) and Bromophenol blue (Cat#B0126, Sigma-Aldrich)) before loading and electrophoresis on 4-20% or 12% SDS-PAGE gel using mini-PROTEAN electrophoresis chamber (Cat#1658001, Bio-Rad). Proteins were transferred onto a 0.45 µm PVDF (Cat#IPVH00010, Millipore) or Nitrocellulose membrane (Cat#GE10600002, GE Healthcare), which was blocked for 30 minutes and incubated with primary antibody in 5% nonfat dry milk in TBST (20 mM Tris, 150 mM NaCl, and 0.05% Tween 20 adjusted to pH 7.6 with 1 M

HCl (Cat#30721, Sigma-Aldrich) overnight at 4°C. The membrane was then incubated with HRP-conjugated secondary antibodies in 5% nonfat dry milk in TBST. Immunoreactive bands were generated by adding Lumi-Light$^{PLUS}$ chemiluminescent substrates (Cat#12015196001, Roche) and chemifluorescence was detected using ChemiDoc MP imaging system (Cat#17001402, Bio-Rad). Western blots presented are representative of at least two independent experiments.

**Metabolic assays**

DHFR activity was detected with DHFR assay kit (Cat#CS0340, Sigma-Aldrich). Mouse and Human tissues were suspended in 200 µl of CelLytic MT reagent (Cat#C3228, Sigma-Aldrich) supplemented with Protease inhibitors and lysed mechanically using glass beads (Cat#G8772, Sigma-Aldrich). Lysates were precleared by centrifugation at 1,500 rpm for 10 minutes. The supernatant was mixed with 6 µl of NADPH, 5 µl of DHFA and 800 µl of Assay buffer provided in the kit in 1 ml quartz cuvette (Cat#104-QS, Hellma) and reaction kinetics were monitored by absorbance at 340 nm for 2-3 minutes using Spectrophotometer. The readouts were normalized to the protein concentration in the supernatant at 280 nm using Nanodrop (Cat#ND-2000, ThermoFisher).

THF levels were analyzed based on a competitive inhibition enzyme immunoassay kit (Cat#CEG411Ge, Cloud-Clone Corp). Dissected E12.5 head samples were suspended in 120 µl of CelLytic MT reagent supplemented with Protease inhibitors followed by mechanical dissociation with a pipette tip for cell lysis and incubated on ice for 30 minutes. Lysates were precleared by ultrasonication and cell debris were pelleted by centrifugation at 1,500 rpm for 10 minutes. The supernatant was separated and kept with kit components at room temperature for 10 minutes. 50 µl of serially diluted standard or samples were added in pre-coated wells provided in the kit followed by incubation and wash steps according to the manufacturer’s instructions. The absorbance was measured at 450 nm using Varioskan flash (Cat#5250030, ThermoFisher) and readouts from standard were plotted to construct a standard curve for estimating THF concentration in head extracts.

SAM levels were determined by Bridge-It assay kit (Cat#1-1-1003, Mediomics) based on biosensor fluorescence. Dissected E12.5 head samples were suspended in 100 µl of supplied CM buffer solution supplemented with Protease inhibitors followed by mechanical dissociation with a pipette tip for cell lysis and incubated at room temperature for 30 minutes. Cell debris were pelleted by centrifugation at 1,500 rpm for 10 minutes and supernatant was separated. In a black 96-well microplate (Cat#237105, ThermoFisher), 10 µl of standard (1 mM SAM) were serially diluted with Buffer S provided in the kit. Then 10 µl of standard mix and 50 µl of samples were incubated with supplied Assay solution for a total volume of 100 µl at room temperature for 30 minutes in dark. The signal intensity of fluorescence excited at 485 nm for 1 second was measured at 665 nm using Varioskan flash. The readouts from standard were plotted to construct a standard curve for estimating SAM concentration in head extracts.

#### QUANTIFICATION AND STATISTICAL ANALYSIS

Fluorescent images were captured with SP8 inverted scanning confocal microscope (Leica Biosystems) or Eclipse 80i fluorescence microscope (Nikon). The microscopic and macroscopic images were prepared with Eclipse TS100 inverted microscope (Nikon) and SMZ18 stereomicroscope (Nikon), respectively. Images were exported as TIFF files and quantification were performed manually with ImageJ (NIH). For experiments involving a pair of conditions, statistical significance between the two sets of data were analyzed with unpaired t test or Mann-Whitney test with Holm-Šidák adjusted p-values using Prism9 (GraphPad). For datasets containing more than two samples, one-way analysis of variance was used to determine adjusted p values. Embryos were collected from different litters, and 100–500 cells were counted for each condition.  Metabolic assays were performed on tissues collected from at least 3 mouse embryos for each genotype. Western blots were performed on tissue lysates collected from 2 embryos for each genotype. Statistically significant differences are reported at $p < 0.05$ (*), $p < 0.01$ (**), $p < 0.001$ (***) and $p < 0.0001$ (****).

##### SEQUENCES

| **Cas9 guide** | **Sequence** |
|---|---|
| 5' | GGATTAGCAACTAGGTAATAAGG |
| 3' | TCCAAGCTGGCCCCATGTTTGGG |

| **Genotyping Primer** | **Sequence** | **Type** |
|---|---|---|
| 1 | GTGATTCCATAGCCTCTTCCAG | Forward |
| 2 | CTTTCCTTTGGTTGCTCTTGTG | Reverse |
| 3 | TAGCACACATGCTCCTTTCC | Reverse |

| **RT-PCR** | **Forward Primer** | **Reverse Primer** |
|---|---|---|
| *Dhfr* (Exon5-6) | CAGGCCACCTCAGACTCTTT | CCTTTTTCCTCCTGGACCTC |

**qRT-PCR**

| **Gene Symbol** | **Forward Primer** | **Reverse Primer** |
|---|---|---|
| *Dhfr* (Exon3-4) | CCATTCCTGAGAAGAATCGACC | CTTTACTTGCCAATTCCGGTTG |
| *Dhfr* (Exon5-6) | CAGGCCACCTCAGACTCTTT | CCTTTTTCCTCCTGGACCTC |
| *Aldoc* | ACCATGACCTCAAACGTTGC | TTGAGCAGAGTCCCTTCGAG |
| *Ttr* | CACCAAATCGTACTGGAAGACA | GTCGTTGGCTGTGAAAACCAC |
| *Gapdh* * | GGCCTTCCGTGTTCCTAC | CTGTTGCTGTAGCCGTATTCA |

* indicates reference/houskeeping gene
